# Potent and reversible open-channel blocker of NMDA receptor derived from dizocilpine with enhanced membrane-to-channel inhibition

**DOI:** 10.1101/2024.04.15.589518

**Authors:** Anna Misiachna, Jan Konecny, Marharyta Kolcheva, Marek Ladislav, Lukas Prchal, Jakub Netolicky, Stepan Kortus, Petra Barackova, Emily Langore, Martin Novak, Katarina Hemelikova, Zuzana Hermanova, Michaela Hrochova, Anezka Pelikanova, Jitka Odvarkova, Jaroslav Pejchal, Jiri Kassa, Jana Zdarova Karasova, Jan Korabecny, Ondrej Soukup, Martin Horak

## Abstract

*N*-methyl-D-aspartate receptors (NMDARs) play a significant role in developing several central nervous system (CNS) disorders. Currently, memantine, used for treating Alzheimer’s disease, and ketamine, known for its anesthetic and antidepressant properties, are two clinically used NMDAR open-channel blockers. However, despite extensive research into NMDAR modulators, many have shown either harmful side effects or inadequate effectiveness. For instance, dizocilpine (MK-801) is recognized for its powerful psychomimetic effects due to its high-affinity and nearly irreversible inhibition of the GluN1/GluN2 NMDAR subtypes. Unlike ketamine, memantine and MK-801 also act through a unique, low-affinity “membrane-to-channel inhibition” (MCI). We aimed to develop an open-channel blocker based on MK-801 with distinct inhibitory characteristics from memantine and MK-801. Our novel compound, K2060, demonstrated effective voltage-dependent inhibition in the micromolar range at key NMDAR subtypes, GluN1/GluN2A and GluN1/GluN2B, even in the presence of Mg^2+^. K2060 showed reversible inhibitory dynamics and a partially trapping open-channel blocking mechanism with a significantly stronger MCI than memantine. Using hippocampal slices, 30 µM K2060 inhibited excitatory postsynaptic currents in CA1 hippocampal neurons by ∼51%, outperforming 30 µM memantine (∼21% inhibition). K2060 exhibited No Observed Adverse Effect Level (NOAEL) of 15 mg/kg upon intraperitoneal administration in mice. Administering K2060 at a 10 mg/kg dosage resulted in brain concentrations of approximately 2 µM, with peak concentrations (Tmax) achieved within 15 minutes. Finally, applying K2060 with trimedoxime and atropine in mice exposed to tabun improved treatment outcomes. These results underscore K2060’s potential as a therapeutic agent for CNS disorders linked to NMDAR dysfunction.

## 1. Introduction

The *N*-methyl-D-aspartate receptors (NMDARs) are ionotropic glutamate receptors crucial in excitatory synaptic transmission in the mammalian central nervous system (CNS). The functional importance of the NMDARs is supported by a large variety of psychiatric and neurological disorders, such as Alzheimer’s disease (AD), autism, intellectual disability, and epilepsy, previously linked with their altered functioning, including those caused by the pathogenic variants within the GluN subunits of the NMDAR (Hu et al., 2016; Strehlow et al., 2019; Garcia-Recio et al., 2020; Hansen et al., 2021). Seven genes encode NMDAR subunits; one gene encodes the GluN1 subunit, four genes encode the GluN2A through GluN2D subunits, and two genes encode the GluN3A and GluN3B subunits (Hansen et al., 2021). The most common conventional NMDARs found in the adult forebrain are diheteromeric GluN1/GluN2A and GluN1/GluN2B, and triheteromeric GluN1/GluN2A/GluN2B receptors (Stroebel et al., 2018; Hansen et al., 2021). All GluN subunits share a typical membrane topology, with an extracellular N-terminal domain, a ligand-binding domain formed by S1 and S2 segments, a transmembrane domain (TMD), and an intracellular C-terminal domain. The TMD consists of three transmembrane-spanning helices (M1, M3, and M4) and one membrane loop (M2) (Wo and Oswald, 1995; Hansen et al., 2021). The M2 loops and M3 helices form the ion channel; its narrowest part is formed by the M2 loops, which function as a selective filter and the binding site for Mg^2+^ at GluN1/GluN2 receptors (Kuner and Schoepfer, 1996; Zhang et al., 2021). Therefore, at resting membrane potentials (for example, -60 mV), the GluN1/GluN2 receptor’s ion channel is blocked by extracellular Mg^2+^ (Nowak et al., 1984; Kuner and Schoepfer, 1996).

There are several critical pharmacological modulators of the NMDARs, for instance, memantine, ketamine, and dizocilpine (MK-801). Despite all being open-channel blockers of the GluN1/GluN2 receptors, memantine is approved for treating AD (Robinson and Keating, 2006), ketamine is used as an anesthetic and antidepressant (Zanos et al., 2018), and MK-801 is used as a model compound mimicking schizophrenia symptoms (Rung et al., 2005). These different pharmacological effects *in vivo* likely result from their differences in the molecular mechanism of action at the NMDARs. Although both memantine and ketamine bind into ion channel pore relatively weakly (”low-affinity” open-channel blockers) (Song et al., 2018; Zhang et al., 2021), memantine—but not ketamine-displays a second type of low-affinity inhibition (Blanpied et al., 1997; Sobolevsky et al., 1998; Kotermanski and Johnson, 2009; Glasgow et al., 2018), by accessing the binding site via a hydrophobic route from the plasma membrane to ion channel through a gated fenestration (named “membrane-to-channel inhibition” (MCI)) (Wilcox et al., 2022). On the contrary, the psychomimetic effect of MK-801 can be ascribed to its “high-affinity “and almost “irreversible” binding within the ion channel pore of GluN1/GluN2 receptors (Song et al., 2018). Recent structural data provided information that the binding site for Mg^2+^ overlaps with the binding site of memantine or ketamine (Song et al., 2018; Zhang et al., 2021). As memantine and ketamine compete with extracellular Mg^2+^ at physiological concentrations (i.e., ∼1 mM), the presence of Mg^2+^ may reduce the increased open-channel blocker’s efficacy (increase the IC_50)_) and may diminish its effect *in vivo* (Kotermanski and Johnson, 2009). Thus, the clinical impact of the open-channel blockers is complex, hardly predictable, and is determined by the distinct pharmacological properties of the individual compound, including selectivity to different subtypes of NMDARs, ability to compete with Mg^2+^, MCI implementation, and the kinetics of onset and offset of inhibition.

Interestingly, also MK-801 exhibits MCI (Wilcox et al., 2022). Here, we developed and characterized, using electrophysiology, a new derivative of MK-801, codenamed K2060, which lacks “irreversible” binding within the ion channel pore of the GluN1/GluN2 receptors while maintaining inhibitory potency even in the presence of 1 mM Mg^2+^ and MCI at clinically relevant concentrations. Given the established benefits of NMDAR antagonists like memantine in mitigating severe seizures induced by organophosphates (Kassa and Karasova, 2021; Gorecki et al., 2022; Kassa and Zdarova Karasova, 2023; Pulkrabkova et al., 2023), our research extended to a mouse model subjected to tabun poisoning. We found that K2060, when used with traditional antidotes such as trimedoxime and atropine, significantly enhanced therapeutic outcomes. Our findings reveal that K2060 possesses unique inhibitory actions at NMDARs, underscoring its promise for future applications in treating CNS disorders linked to dysregulated NMDAR activity.

## 2. Materials and methods

### 2.1. General Chemistry

All chemical solvents and reagents were used in the highest available purity without further purification. They were purchased from Sigma-Aldrich (Prague, Czech Republic) or FluoroChem (Dublin, Ireland). The reactions were monitored by thin layer chromatography on silica gel plates (60 F254, Merck, Prague, Czech Republic), and the spots were visualized by ultraviolet light (254 nm). Purification of crude product was carried out using PuriFlash GEN5 column, 5.250 (Interchim, Montluçon, France; silica gel 100, 60 Å, 230– 400 mesh ASTM, Sigma-Aldrich). NMR spectra were recorded in deuterated methanol (CD_3_OD) on Bruker Avance NEO 500 MHz spectrometer (499.87 MHz for ^1^H NMR and 125.71 MHz for ^13^C NMR). Chemical shifts (*δ*) are reported in parts per million (ppm), and spin multiplicities are given as broad singlet (bs), doublet (d), doublet of doublet (dd), triplet (t), quartet (q), pentet (p), or multiplet (m). Coupling constants (*J*) are reported in Hz. The final compound was analyzed by an LC-MS system consisting of UHLPC Dionex Ultimate 3000 RS coupled with a Q Exactive Plus Orbitrap mass spectrometer to obtain high-resolution mass spectra (Thermo Fisher Scientific, Bremen, Germany). The melting point was measured using an automated melting point recorder M-565 (Büchi, Switzerland) and is uncalibrated. Gradient LC analysis confirmed > 99% purity for the final compound.

### 2.2. Chemical Procedure for K2060

Commercially available 5*H*-dibenzo[*a,d*][7]annulen-5-ol (**1**; 4.8 mmol, 1.0 eq; Fig. 1) was placed into a 25 mL flask and dissolved in dry benzene; the flask was filled with argon. Solution of SOCl_2_ (20.67 mmol, 4.3 eq), 1 mL of dry benzene, and 0.1 mL of pyridine was added dropwise at room temperature. The reaction mixture was stirred at 85 °C for 2 hours under an argon atmosphere. After the reaction proceeded, the solvent was evaporated under reduced pressure, and the crude material was crystallized from petroleum ether to obtain the final compound (**2**; Fig. 1) as a yellow solid. Intermediate **2** was used directly for the following reaction. Cyclobutylamine (**3**; 5.25 mmol, 3 eq) was placed in a 50 mL flask and dissolved in 10 mL of dry MeCN. The reaction was kept under an argon atmosphere at room temperature. 5-Chloro-5*H*-dibenzo[*a,d*][7]annulene (**2**; 1.75 mmol, 1.0 eq) was dissolved in 5 mL of dry MeCN and added dropwise into the reaction. The reaction was stirred for 3 hours under an argon atmosphere at 40 °C. After cooling to room temperature, the solvent was evaporated under reduced pressure, and the crude material was purified using flash chromatography (eluent CH_2_Cl_2_/MeOH with 1% of 25% aq. NH_3_, gradient elution 98:2 → 94:6) to obtain the final compound as a free base. This was dissolved in 5 mL of MeOH, and 1 mL of HCl (35% aq. sol.) was added dropwise at 0 °C. The reaction was maintained stirring for 1 hour at room temperature. The solvent was evaporated under reduced pressure, and the residual water was removed by azeotropic distillation with absolute EtOH. The solid product was washed with cold acetone, and a white solid of K2060 hydrochloride salt was collected by filtration.

**Fig. 1.**
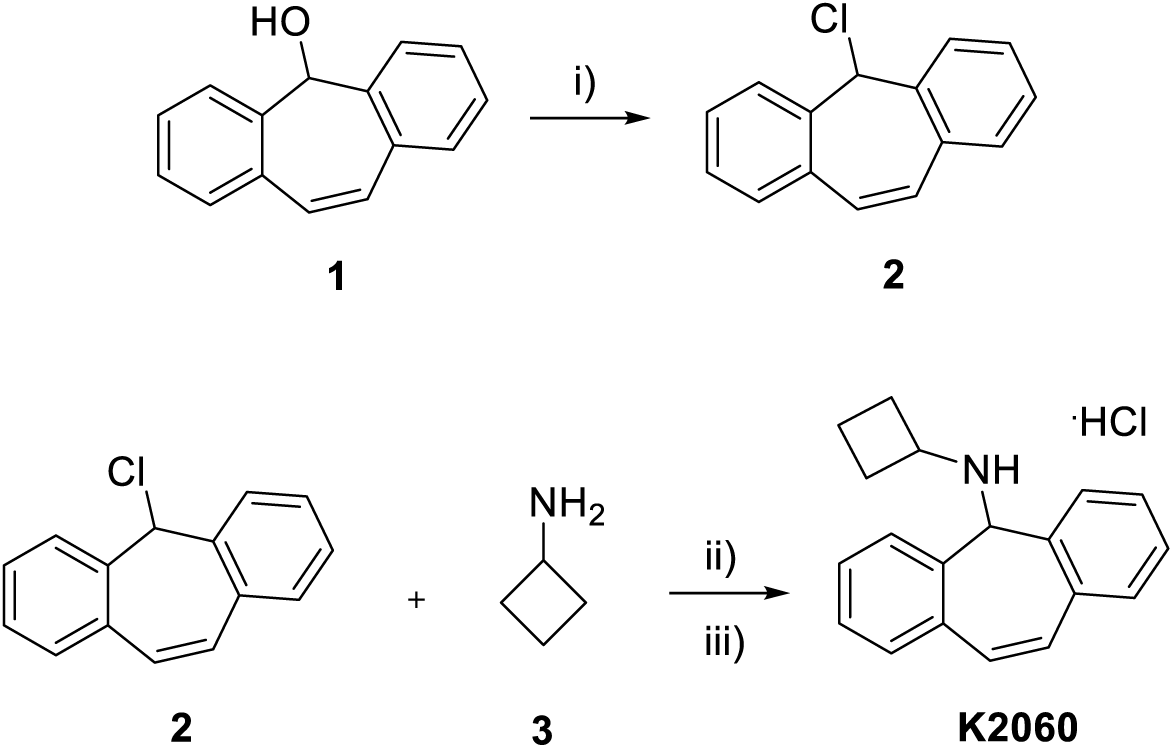
Synthetic procedure for the preparation of K2060. Reaction conditions: (i) SOCl_2_, benzene, 85 °C, 2 h; (ii) MeCN, 40 °C, 3 h and (iii) HCl (37% aq.sol.), MeOH, RT, 1 h.

### 2.3. N-Cyclobutyl-5H-dibenzo[a,d][7]annulen-5-amine hydrochloride (K2060)

Total yield 64 %; white solid. M.p.: 217.7 – 219.1 °C. ^1^H NMR (500 MHz, methanol-*d*_4_) *δ* 7.74 – 7.69 (m, 2H), 7.63 – 7.59 (m, 2H), 7.58 – 7.52 (m, 4H), 7.25 (s, 2H), 5.63 (s, 1H), 3.40 – 3.34 (m, 1H), 2.15 – 2.03 (m, 2H), 2.02 – 1.92 (m, 2H), 1.83 – 1.64 (m, 2H). ^13^C NMR (126 MHz, MeOD) *δ* 134.03, 130.94, 130.51, 130.21, 130.13, 129.47, 129.43, 65.83, 50.94, 26.44, 14.42. HRMS (ESI^+^): [M + H]^+^: calculated for C_19_H_20_N^+^ (m/z): 262.15902; found: 262.15839. LC-MS purity 99.9%.

### 2.4. Molecular biology and HEK293 cell culture

The following cDNA vectors encoding the full-length human GluN subunits were employed: human GluN1-4a (GluN1), human GluN2A (GluN2A), and human GluN2B (GluN2B), the reason to use the GluN1-4a splice variant is that it produces larger NMDAR current responses than the GluN1-1a splice variant (Kolcheva et al., 2021). The Quick-Change site-directed mutagenesis kit (Agilent, Santa Clara, CA) was used to generate the specific point mutations within the coding regions of the GluN subunits. Human embryonic kidney 293 (HEK293) cells were grown in Opti-MEM I media containing 5% (v/v) fetal bovine serum (FBS; Thermo Fischer Scientific). HEK293 cells grown in 24-well plates were transfected in 50 μl Opti-MEM I with 0.9 μl MATra-A Reagent (IBA, Louvain-La-Neuve, Belgium) and a total of 0.9 μg of cDNA vectors encoding the GluN1/GluN2 receptors and GFP (to identify the transfected cells; pQBI 25, Takara, Shiga, Japan). After transfection, the HEK293 cells were trypsinized and grown at a lower density on poly-L-lysine‒coated glass coverslips (diameter 3 cm) in Opti-MEM I containing 1% (v/v) FBS, 2 mM MgCl_2_, 1 mM DL-2-amino-5-phosphonovaleric acid, and 3 mM kynurenic acid (to reduce excitotoxicity induced by the activation of the NMDARs).

### 2.5. Electrophysiology in HEK293 cells

Whole-cell patch-clamp recordings were performed from the transfected HEK293 cells ∼16-48 h after the transfection using an Axopatch 200B amplifier (Molecular Devices, San Jose, CA, USA) with series resistance (<10 MΩ) and ∼80% capacitance compensation at a holding membrane potential of -60 mV (unless stated otherwise). The current responses were low-pass filtered at 2 kHz using an eight-pole Bessel filter, digitized at 5 kHz, and recorded using pCLAMP 11 (Molecular Devices). Borosilicate glass pipettes with a tip resistance of 3–6 MΩ were prepared using a P-97 puller (Sutter Instruments, Novato, CA, USA) and filled with intracellular recording solution (ICS) containing (in mM): 125 gluconic acid, 15 CsCl, 5 EGTA, 10 HEPES, 3 MgCl_2_, 0.5 CaCl_2_, and 2 ATP-Mg salt (pH 7.2 with CsOH). The extracellular recording solution (ECS) contained (in mM) 160 NaCl, 2.5 KCl, 10 HEPES, 10 D-glucose, 0.01 EDTA, and 1 CaCl_2_ (pH 7.3 with NaOH). The ECS contained a saturating concentration of 50 µM glycine, which is not indicated in the figures. The NMDAR current responses were induced by a saturating concentration of 1 mM L-glutamate in the ECS, using a multi-barrel rapid application system with a time constant of solution exchange around the measured cell of ∼20 ms (Kolcheva et al., 2021). EDTA was omitted from the ECS for measuring in the presence of extracellular 1 MgCl_2_. The stock solution of K2060 (10 mM) was prepared in dimethyl sulfoxide (DMSO) and stored at -20 °C for several months. All electrophysiological recordings were performed at room temperature (RT), and the reported membrane potential values were not corrected for the liquid junction potential.

The concentration-response curves for K2060 were obtained using the following equation:

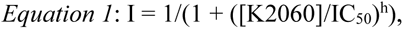

IC_50_ is the concentration of K2060 that produced 50% inhibition of the agonist-evoked current, [K2060] is the concentration of K2060, and h is the apparent Hill coefficient. The time course of K2060 inhibition was analyzed using a double exponential function, and weighted time constants (τ_w_) were obtained using the following equation:

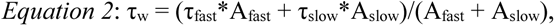

The faster component has a time constant of τ_fast_ and amplitude A_fast_, and the slower component has a time constant of τ_slow_ and amplitude A_slow_; normalized values of A_fast_ were obtained by dividing each value by (A_fast_ + A_slow_)(Glasgow et al., 2018). MCI inhibition was measured as described (Glasgow et al., 2018). The minimum fractional response after MCI (minimum I_MCI_/I_control_) was calculated as the mean value of I_MCI_/I_control_ measured over a 200-ms window centered on the minimum ratio value; I_control_ was calculated as the mean of I_control1_ and I_control2_; the responses in which I_control2_/I_control1_ was <0.8 were excluded from analysis (Glasgow et al., 2018). The recovery time constant from MCI (τ_recovery_) was obtained by fitting the I_MCI_/I_control_ values with a single-exponential function.

### 2.6. Electrophysiology in CA1 pyramidal neurons

Recordings were conducted in hippocampal slices obtained from young adult WT C57BL/6N mice of both sexes (P30-35). The mice were deeply anesthetized with inhaled isoflurane, decapitated, and their brains were swiftly removed and submerged in ice-cold oxygenated (95% O_2_ / 5% CO_2_) artificial cerebrospinal fluid (aCSF) with the following composition (in mM): 124 NaCl, 3.5 KCl, 1.25 NaH_2_PO_4_, 26 NaHCO_3_, 4 MgSO_4_, 0.5 CaCl_2_, and 11 D-glucose (pH 7.3). Horizontal 300 μm slices were cut in an ice-cold oxygenated aCSF bath; the slices were hemisected and then incubated for 30 minutes in oxygenated aCSF with 1 mM MgSO_4_ and 1 mM CaCl_2_ at 32°C. Subsequently, the slices were equilibrated to RT for 30 minutes and kept at RT for at least 60 minutes or until used for recordings. Recordings were performed from hippocampal CA1 pyramidal neurons using a Multiclamp 700B amplifier (Molecular Devices), sampled at 25 kHz (Digidata 1550B, Molecular Devices), and low-pass filtered at 10 kHz using pCLAMP 11. Neurons were visualized by Dodt gradient contrast with 40x water-immersion objective and a digital video camera (Axiocam 305 color, Zeiss, Prague, Czech Republic) using AxioExaminer.Z1 microscope (Zeiss). The recording aCSF (300-310 mOsm/kg) was identical to holding aCSF and was maintained at 32°C for all recordings. Slices were selected randomly for recording, and one cell was recorded per slice. For voltage-clamp recordings, borosilicate glass recording electrodes (4-5 MΩ) were filled with (in mM) 120 Cs-Gluconate, 9 NaCl, 10 HEPES, 10 BAPTA, 4 ATP-Mg, 0.3 GTP-Mg, 2 MgCl_2_ (pH 7.3 adjusted by CsOH, osmolarity adjusted to 290 mOsm/Kg). Following a 10-minute equilibration period, evoked excitatory postsynaptic currents (eEPSCs) at a holding potential -60 mV (liquid junction potential was not corrected) were recorded every 20 seconds using stimulus generator STG5 (Multichannel systems, Reutlingen, Germany) triggered by Clampex 11. To evoke eEPSCs, a concentric bipolar tungsten electrode (World Precision Instruments, Sarasota, FL, USA, #TM33CCINS-B) was placed to the stratum radiatum, and a bipolar square pulse for 100 µs/phase was applied. To isolate the NMDAR component of eEPSCs, 10 µM NBQX (inhibitor of α-amino-3-hydroxy-5-methyl-4-isoxazole propionic acid receptor; AMPAR) and 100 µM picrotoxin (inhibitor of gamma-aminobutyric acid receptor; GABAR) were applied during whole experiments. The train of 5 stimuli at 50 Hz (20 ms interstimulus interval) was applied to assess the inhibitory effect of K2060 and memantine (Hello Bio, Bristol, UK). The stability of trains was ensured by a 5 – 15-minute stability period before the K2060 or memantine was applied; trains were applied at the beginning and the end of this period. Four trains were recorded before and after 10 min application of drug/vehicle; then they were averaged and analyzed using Clampfit 11. The current from the fifth stimulus was used to assess inhibitory effects. Series resistance and cell capacitance were continually monitored but remained uncompensated throughout the experiment. Cells were excluded from the analysis if either parameter exhibited a change exceeding 20% during the experiment.

### 2.7. Animals

Adult male and female ICR mice (30-40 g) purchased from the VELAZ (Prague, Czech Republic) were used for all *in vivo* studies. The animals were kept in an air-conditioned enclosure with light from 07:00 *a.m.* to 7:00 *p.m.* and were allowed access to standard food and tap water *ad libitum*. Mice were divided into groups of 4 (toxicity study, pharmacokinetic study) and 6 animals (overall protection study), respectively. The mice were acclimatized in the laboratory vivarium for 14 days before the experiments commenced. The compound K2060 was administered intraperitoneally (i.p., 5% DMSO with 5% Kolliphor in saline in 0.1 ml/kg) in acute toxicity evaluation and pharmacokinetic study, and intramuscularly (i.m., 5% DMSO with 5% Kolliphor in saline in 0.1 ml/kg) in overall protection study. All *in vivo* studies followed ethical guidelines: The guidelines of the European Union conducted the experiments directive 2010/63/E.U. and Act No. 246/1992 Coll. The experimental animals were handled under the supervision of the Ethics Committee of the Military Faculty of Medicine, Czech Republic (approval reference No. MO149149/2022-1457, MO127213/2024-1457).

### 2.8. Acute toxicity evaluation-NOAEL

Adult male and female ICR mice were used to assess the no-observed-adverse-effect-level (NOAEL) of K2060 as the acute toxicity marker. Mice were randomly assigned to the experimental groups consisting of four males and four females per one administered dose of K2060. Selected doses were administered to identify NOAEL with starting dose levels based on their solubility or according to our previous experience. Treated mice were extensively observed for signs of toxicity in the first hour and then periodically for two days. Clinical signs such as cardiovascular, respiratory, and nervous system disability, weight loss, or reduction of food consumption were observed according to Laboratory Animal Science Association (U.K.) guidelines. The severity of symptoms was classified as mild, moderate, and substantial (Robinson et al, 2009). During all acute toxicity studies, we followed the rules: **i)** if the category of substantial severity was achieved, the animal was immediately euthanized by CO_2_, and a lower/more appropriate dose was selected to continue the study; **ii)** if a severe adverse effect or death occurred within a few minutes after administration to the first animal in the group, the other animal was not treated, and a lower dose was given; **iii)** in the case of mortality or surviving 48 hours, all animals were euthanized by CO_2_ and subjected to basic macroscopic necropsy (Misik et al., 2018).

### 2.9. In vivo pharmacokinetic study

Adult male ICR mice were used for the pharmacokinetic assessment of K2060. K2060 was administered at 10 mg/kg (i.e., just below the NOAEL dose). Blood samples were collected under deep terminal anesthesia directly by cardiac puncture into heparinized 1.5 mL tubes at 5, 10, 15, 20, 30, 60, 120, 240, and 360 min (four animals per time interval). Four animals were used for zero time/blank. The animals were perfused transcardially with a saline solution (0.9% NaCl) for 5 min (1 mL/min), and after the washout, the skull was opened, and the brain was carefully removed; brains were stored at −80 °C until homogenization. The brains were weighed, and four times the weight of phosphate-buffered saline (PBS) was added. The brains were subsequently homogenized by a T-25 Ultra Turrax disperser (I.K.A., Staufen, Germany), ultrasonicated by UP 50H needle homogenizer (Hielscher, Teltow, Germany), and stored at -80 °C before extraction. 190 µL of brain homogenate or 95 µL of plasma were spiked with 10 µL (5 µL for plasma samples) of internal standard (IS, donepezil in methanol, final concentration was 100 nM). The sample was alkalized with 100 µL (50 µL for plasma samples) of 1 M sodium hydroxide, and 700 µL (350 µL for plasma samples) of tert-butyl methyl ether was added. The samples were then shaken (1200 RPM, 10 min, Wizard Advanced I.R. Vortex Mixer, Velp Scientifica, Usmate, Italy) and centrifuged (12,000 RPM, 5 min, Universal 320 R centrifuge, Hettich, Tuttlingen, Germany). 500 µL (250 µL for plasma samples) of supernatant was transferred to a microtube and evaporated to dryness in a CentriVap concentrator (Labconco Corporation, Kansas City, USA). Calibration samples were prepared by the spiking of 180 µL of blank brain homogenate or 95 µL of plasma with 10 µL (5 µL for plasma samples) of the studied compound dissolved in methanol (final concentrations ranged from 0.5 nM to 50 µM) and with 10 µL (5µL) of IS, with a final concentration of 100 nM. The calibration samples were then vortexed and extracted as above. The analyzed samples were reconstituted in 100 µL (50 µL µL for plasma samples) of acetonitrile/water mixture of 50/50 (v/v).

The UHPLC (mentioned above) system with mass spectrometric detection measured compound levels in plasma and brain homogenate. The results were obtained by the gradient elution method with a reverse phase on an RP Amide column (Ascentis Express, 2.1x100 mm, 2.7 µm, Merck, Darmstadt, Germany). Mobile phase A was ultrapure water of ASTM I type (resistance 18.2 MΩ.cm at 25°C) prepared by Barnstead Smart2Pure 3 UV/UF apparatus (Thermo Fisher Scientific, Bremen, Germany) with 0.1 % (v/v) formic acid (LC-MS grade, VWR International, Radnor, PA, USA); mobile phase B was acetonitrile (MS grade, VWR International, Radnor, PA, USA) with 0.1 % (v/v) of formic acid. The gradient was as follows. Initially, 10 % of mobile phase B flowed for 0.3 min; the gradient of mobile phase B then increased to 60 % during 2.7 min and then rose to 100 % in 0.7 min. After this 0.7 min of 100 % B flow, the composition returned to 10 % B and was equilibrated for 3.5 min. The total run time of the gradient elution method was 7.5 min. The mobile phase flow rate was set to 0.45 mL/min, and the column was tempered to 45 °C. The injection volume was 5 µL. Samples were analyzed using the previously mentioned Q Exactive Orbitrap mass spectrometer set to collect the total ion current in positive mode. The calibration curves had 9 points, and concentrations ranged from 5 nM to 50 µM; curves in both cases were linear within the measured range.

### 2.10. In vivo protection study

Adult male ICR mice were used for the protection study; for each sex, they were divided into groups of six animals. Tabun was obtained from the Military Technical Institute (Brno, Czech Republic); before the experiment, the purity was verified by gas chromatography-mass spectrophotometry and exceeded 95%. The tabun stock solution (1 mg/mL) was prepared in propylene glycol 3 days before starting the experiments. The solution was diluted with saline immediately before administration. Standard oxime reactivator trimedoxime was synthesized at the Department of Toxicology and Military Pharmacy, the Military Faculty of Medicine in Hradec Kralove. All other similar analytical-grade chemicals were purchased commercially and used as received. All compounds were administered by i.m. injection at a volume of 10 mL/kg body weight (b.w.). K2060 was administered at 10 mg/kg (i.e., just below the NOAEL dose). Trimedoxime was administered at a dose corresponding to 5% of its median lethal dose (LD_50_; 5.3 mg/kg) in combination with atropine (10 mg/kg). The therapeutic efficacy of K2060, classic antidotal treatment, or their combination was studied when antidotes were applied 1 min after tabun intoxication. NA-induced toxicity was determined by assessing LD_50_ value and its 95% confidence limit using probit-logarithmical analysis of death occurring within 24 h after administration of N.A.s at five different doses with six animals per dose (Tallarida et Murray, 1986). The therapeutic efficacy of all antidotes was expressed as protective ratio A (LD_50_ value of NA in treated mice/LD_50_ value of NA in non-treated mice) and protective ratio B (LD_50_ value of NA in treated mice with the combination of K2060 with classic antidotal treatment/LD_50_ value of NA in treated mice with classic antidotal treatment).

### 2.11. Statistical analyses

Unless stated differently, summary data are presented as the mean ± the standard error of the mean (S.E.M.); group differences were analyzed using the Student’s *t*-test or one-way ANOVA followed by Tukey’s post hoc test using SigmaStat 3.5 (Systat Software Inc.). Differences with a *p*-value < 0.05 were considered significant.

## 3. Results

### 3.1. Chemical Synthesis

A long-term search for novel modulators of NMDARs identified a hit compound (**K2060**) synthesized from commercially available 5*H*-dibenzo[*a*,*d*][7]annulen-5-ol (**1**). The starting material was converted by nucleophilic substitution to 5-chloro-5*H*-dibenzo[*a*,*d*][7]annulene (**2**; Fig. 1). Reaction with cyclobutylamine (**3**) allowed the formation of amine, which was subsequently converted to a hydrochloride salt (**K2060**).

### 3.2. The inhibitory effect of K2060 at GluN1/GluN2A and GluN1/GluN2B receptors

First, we measured the inhibitory concentration-response curves for K2060 (using a range of 0.3-300 μM) at GluN1/GluN2A and GluN1/GluN2B receptors activated with 1 mM L-glutamate at membrane potentials of -60 and +40 mV, both in the absence and presence of 1 mM Mg^2+^ (Fig 2A-D). The resulting IC_50_ values for K2060 and, for comparison, the IC_50_ values measured under the same conditions for memantine are shown in Table 1. Regarding the GluN1/GluN2A receptors, our experiments showed that i.) K2060 acts as a voltage-dependent inhibitor (as does memantine), ii.) when measured without extracellular Mg^2+^, K2060 has an IC_50_ value ∼2.0 times smaller than memantine at a membrane potential of -60 mV but a similar IC_50_ value at a membrane potential of +40 mV, and iii.) K2060 is ∼2.7 times more potent at a membrane potential of -60 mV than memantine in 1 mM Mg^2+^ (Table 1). For GluN1/GluN2B receptors, we observed that K2060 i.) also acts as a voltage-dependent inhibitor (as memantine), ii.) when measured without extracellular Mg^2+^, K2060 has slightly lower IC_50_ values than memantine at membrane potentials of -60 mV and +40 mV, and iii.) K2060 is ∼2.2 times more potent at a membrane potential of -60 mV than memantine in 1 mM Mg^2+^. Comparing the inhibitory effect of K2060 between GluN1/GluN2A and GluN1/GluN2B receptors, we observed similar IC_50_ values at a membrane potential of -60 mV and ∼2.3-fold lower IC_50_ value for GluN1/GluN2B receptor at a membrane potential of +40 mV, both at 0 and 1 mM Mg^2+^. These results showed that K2060 did not exhibit a subunit-dependent effect in the studied NMDARs.

**Fig. 2.**
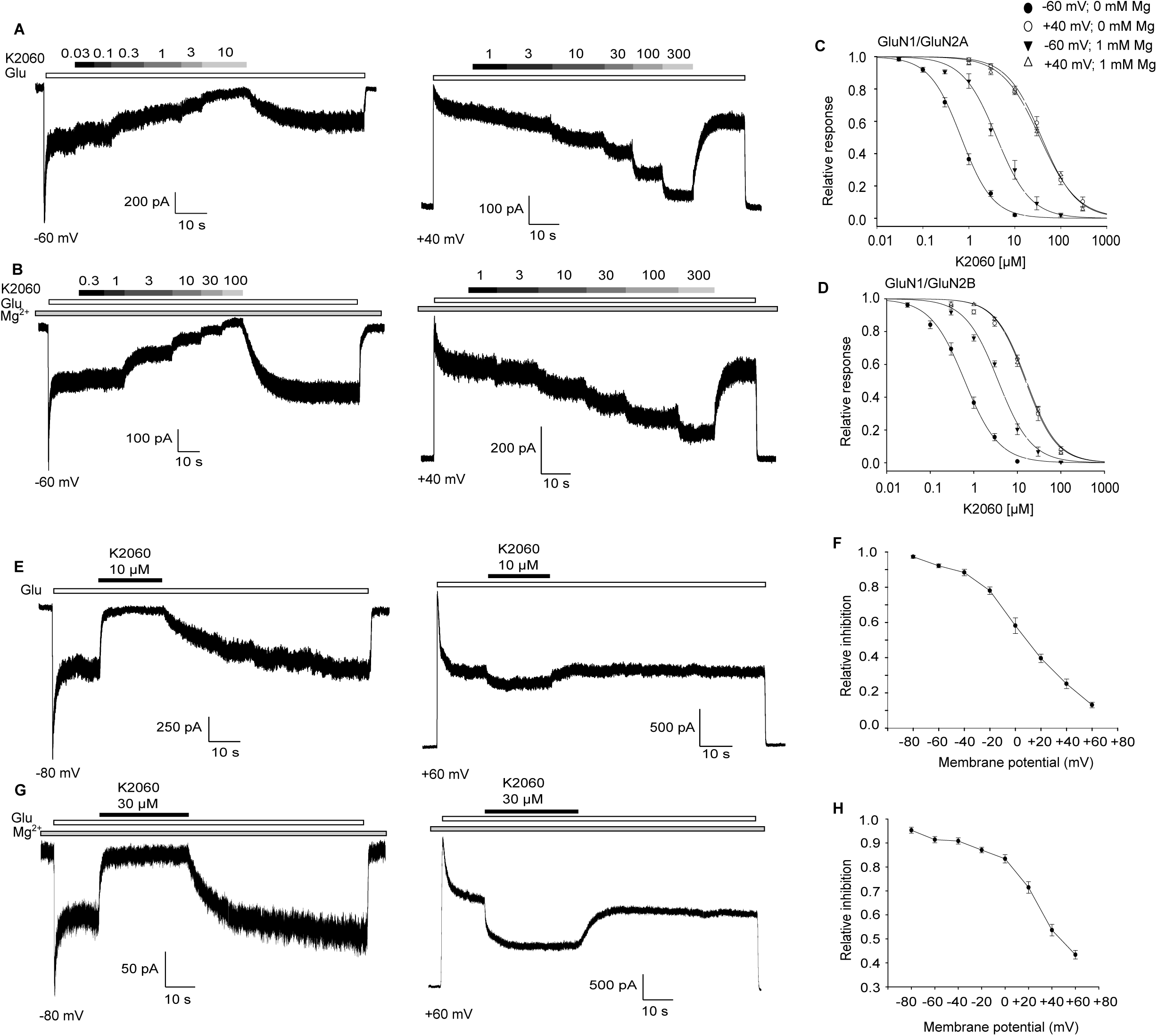
K2060 is a potent voltage-dependent inhibitor of GluN1/GluN2 receptors. (A, B) Representative current traces recorded at indicated membrane potentials from HEK293 cells transfected with GluN1/GluN2A receptors. The indicated concentrations of K2060 (in µM) were applied in the absence (A) or the continuous presence (B) of 1 mM Mg^2+^. (C, D) Concentration-response curves for inhibition by K2060 were obtained by fitting the experimental data from HEK293 cells transfected with the GluN1/GluN2A (C) and GluN1/GluN2B (D) receptors. The IC_50_ values obtained by fitting the electrophysiological data using Equation 1 (see Methods) are shown in Table 1. (E, G) Representative current traces from HEK293 cells transfected with GluN1/GluN2A receptors showing the inhibitory effect of 10 μM (E) or 30 μM (G; in the presence of 1 mM Mg^2+^) K2060 applied at the indicated membrane potentials. (F, H) The graphs showing the relative inhibition of the current responses of the GluN1/GluN2A receptor, induced by 10 μM (F) or 30 μM (H; in the presence of 1 mM Mg^2+^) K2060 at membrane potentials spanning from -80 to +60 mV at 20 mV steps.

**Table 1.** The IC_50_ values for inhibitory effects of memantine (Mem) and K2060 at the NMDARs.

| Receptor | Compound | 0 mM $Mg^{2+}$<br>$IC_{50} \pm S.E.M., \mu M$ | | 1 mM $Mg^{2+}$<br>$IC_{50} \pm S.E.M., \mu M$ | |
| --- | --- | --- | --- | --- | --- |
|  |  | -60 mV | +40 mV | -60 mV | +40 mV |
| GluN1/GluN2A | Mem | $1.34 \pm 0.08$ (n=6) | $39.84 \pm 1.41$ (n=5) | $11.75 \pm 0.53$<br>(n=7) | $36.31 \pm 1.78$<br>(n=10) |
| | K2060 | $0.68 \pm 0.07$ (n=7) | $38.30 \pm 3.70$ (n=7) | $4.40 \pm 0.62$ (n=5) | $35.13 \pm 3.10$ (n=6) |
| GluN1/GluN2B | Mem | $0.78 \pm 0.09$ (n=5) | $25.63 \pm 1.36$ (n=7) | $8.05 \pm 0.42$ (n=5) | $22.67 \pm 1.84$ (n=4) |
| | K2060 | $0.62 \pm 0.07$ (n=7) | $16.11 \pm 1.59$<br>(n=14) | $3.62 \pm 0.09$ (n=5) | $15.35 \pm 1.05$ (n=8) |

Next, we focused on studying the mechanism of K2060’s inhibitory effect at the GluN1/GluN2A receptor. It generates larger current responses in HEK293 cells than the GluN1/GluN2B receptor, simplifying electrophysiological studies in the presence of 1 mM Mg^2+.^ To verify that K2060 acts as a voltage-dependent inhibitor over the entire range of membrane potentials, we further recorded the relative inhibition rate of 10 μM (in the case of 0 mM Mg^2+^) or 30 μM (in the case of 1 mM Mg^2+^) K2060 at membrane potentials from -80 mV to +60 mV (in 20 mV steps; Fig. 2E,G). These experiments showed that K2060 inhibited the GluN1/GluN2A receptor over the entire range of measured membrane potentials but more strongly at negative membrane potentials (Fig. 2F,H). These electrophysiological experiments showed that K2060 is a potent voltage-dependent inhibitor of GluN1/GluN2A and GluN1/GluN2B receptors even in the presence of 1 mM Mg^2+^ and thus likely acts as an open-channel blocker.

A critical parameter that regulates the pharmacological efficacy of open-channel blockers of NMDARs is the rate of their inhibitory effect (i.e., onset and offset kinetics). Interestingly, Mg^2+^ accelerates the offset kinetics by memantine (Glasgow et al., 2018). We next studied the rate of onset and offset of inhibition mediated by 3 or 10 µM K2060 at the GluN1/GluN2A receptor, both in the absence and presence of 1 mM Mg^2+^ (Fig. 3A). Our measurements showed that the τ_on_ values of the inhibitory effect of K2060 were ∼1.2 s for 3 µM K2060 and ∼0.4 s for 10 µM K2060 in the absence of Mg^2+^. In contrast, in the presence of 1 mM Mg^2+^, τ_on_ values increased to ∼2.7 s for 3 µM K2060 and ∼1.4 s for 10 µM K2060 (Fig. 3B). The measured τ_off_ values were in the range of ∼9 s for both 3 and 10 µM K2060 in the absence of Mg^2+^, while in the presence of 1 mM Mg^2+^, they decreased to ∼4 s for both 3 µM and 10 µM K2060 (Fig. 3C). Compared to our previous data for 10 µM memantine (τ_on_ - 0 mM Mg^2+^: ∼100 ms, 1 mM Mg^2+^: ∼200 ms; τ_off_ - 0 mM Mg^2+^: ∼800 ms, 1 mM Mg^2+^: ∼400 ms) (Kolcheva et al., 2023), our experiments showed that K2060 inhibits GluN1/GluN2A receptor with slower kinetics compared to memantine. Because the open-channel blockers of NMDARs can act through MCI (see below), we next asked whether different application times of K2060 alter its offset kinetics (Fig. 3D). These measurements revealed no difference in the measured τ_off_ values after the application times of K2060 in the range of 3, 10, 30, and 100 s (Fig. 3E), confirming that K2060 unbinds from the GluN1/GluN2A receptor with slow kinetics independent of the time of application of K2060. The open-channel blockers of NMDARs can act by a “foot-in-the-door” mechanism (they block only the open state of the receptor and do not allow the release of the agonist), a “trapping” mechanism (they block the ion channel in a way that enables the release of the agonist, the blocker is released in the presence of the agonist), or a “partial trapping” mechanism (they allow some of the blocker to escape from the ion channel even in the absence of the agonist) (Sobolevsky et al., 2002; Vyklicky et al., 2015). To determine the blocking mechanism of K2060, we next asked whether K2060 unbinds from the GluN1/GluN2A receptor in the absence of L-glutamate. We co-applied 1 mM L-glutamate and 10 µM K2060 until a steady state of K2060-mediated inhibition has been reached; then, we applied ECS (without L-glutamate and K2060) for 10 s or 100 s and then activated GluN1/GluN2A receptor using 1 mM L-glutamate (Fig. 3F). We chose 10 s of ECS application after inhibition of the GluN1/GluN2A receptor with K2060 because our MCI measurements showed that K2060 is almost wholly removed from the membrane within 10 s (described below), and for comparison, one order of magnitude higher time, i.e., 100 s. This experiment showed that the relative inhibition by K2060, measured 200 ms after initiation of L-glutamate application, is significantly reduced after 100 s of ECS application (∼0.3) compared to 10 s of ECS application (∼0.2; Fig. 3G,H). Therefore, this experiment ruled out a “trapping” mechanism for the blocking effect of K2060 at GluN1/GluN2A receptors. In the following experiment, we observed that the inhibitory effect of 10 µM K2060 is completely abolished within a 100 s period in the presence of 1 mM L-glutamate (Fig. 3I; consistently with measured τ_off_ values), thus ruling out a “foot-in-the-door” mechanism (because these electrophysiological measurements took several minutes, the last two experiments were not performed on the same cell). Previously, we analyzed “tail currents” after the removal of L-glutamate and 7*-*methoxy derivative of tacrine (7-MEOTA) to analyze the mechanism of its blocking effect (Kaniakova et al., 2018). Thus, we further investigated whether K2060 produces “tail currents,” so we simultaneously applied 1 mM L-glutamate and K2060 until its inhibitory effect reached a steady state; then, we removed both L-glutamate and K2060 (Fig. 3J). We did not observe any “tail currents” at GluN1/GluN2A receptors when 10 or 30 µM K2060 was used, precluding the possibility of studying the mechanism of blockade by K2060 using “tail current” analysis. Our electrophysiological data indicate that K2060 acts by a “partial trapping” mechanism at the GluN1/GluN2A receptor.

**Fig. 3.**
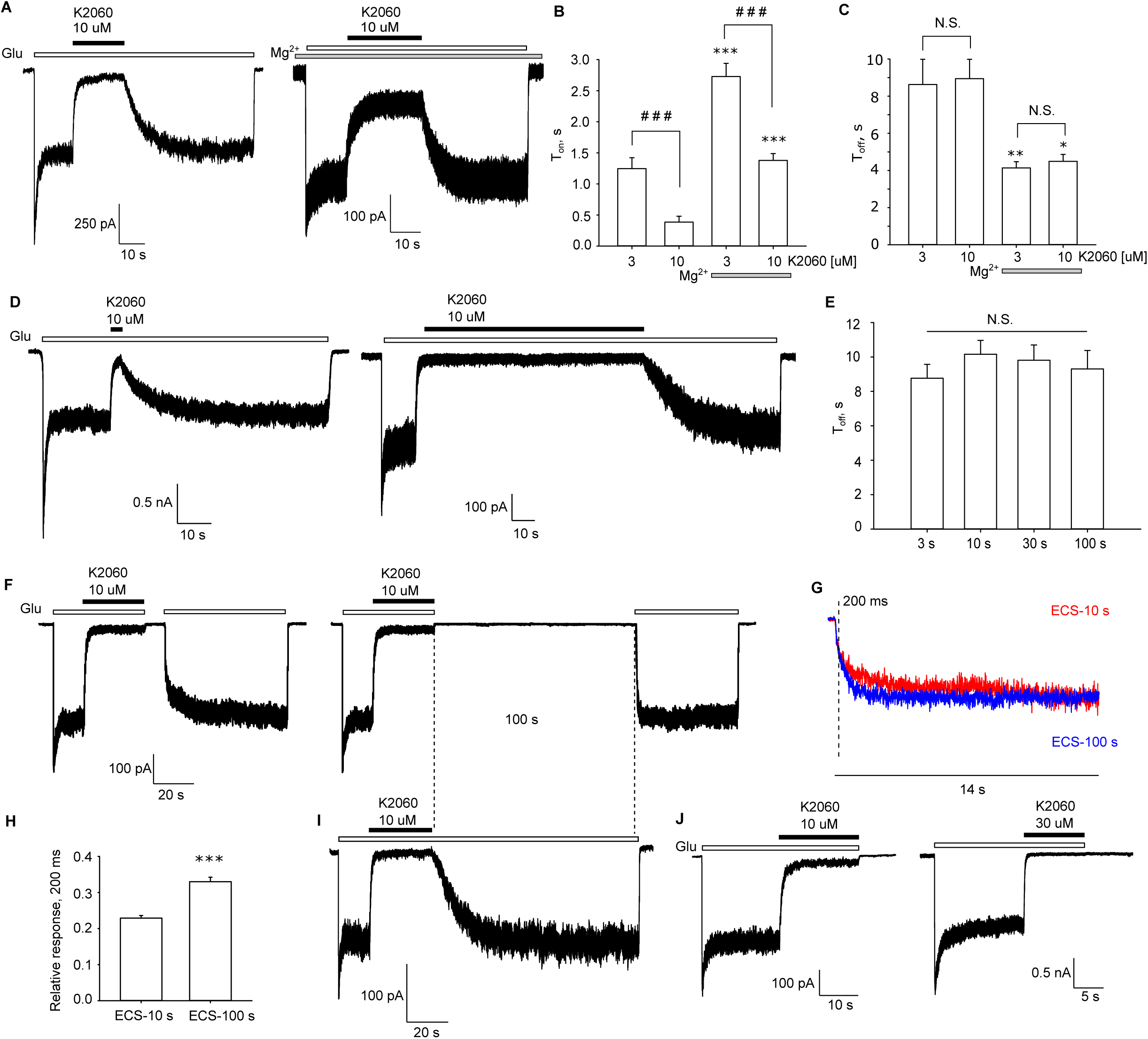
K2060 likely acts as a “partially trapping” open-channel blocker of the GluN1/GluN2A receptor. (A) Representative current traces from HEK293 cells recorded at a membrane potential of -60 mV showing inhibitory kinetics of 10 μM K2060 at GluN1/GluN2A receptor in the absence or the presence of 1 mM Mg^2+^. (B, C) Weighted time constants of the onset (B, τ_on_) and offset (C, τ_off_) inhibitory kinetics of K2060. Time constants were obtained by fitting the experimental data with a double exponential function using Equation 2 (t_(15)_ = -5.537, ***p = <0.001, Student’s t-test (τ_on_ 3 µM versus 3 µM+Mg^2+^); t_(14)_ = - 11.389, ***p = <0.001, Student’s t-test (τ_on_ 10 µM versus 10 µM+Mg^2+^); t_(18)_ = 6.065, ^###^p = <0.001, Student’s t-test (τ_on_ 3 µM versus 10 µM); t_(11)_ = 4.718, ^###^p = <0.001, Student’s t-test (τ_on_ 3 µM+Mg^2+^ versus 10 µM+Mg^2+^); t_(13)_ = 3.410, **p = 0.005, Student’s t-test (τ_off_ 3 µM versus 3 µM+Mg^2+^); t_(12)_ = 3.067, *p = 0.010, Student’s t-test (τ_off_ 10 µM versus 10 µM+Mg^2+^); t_(14)_ = -0.187, p = 0.855, Student’s t-test (τ_off_ 3 µM versus 10 µM); t_(11)_ = -0.688, p = 0.505, Student’s t-test (τ_off_ 3 µM+Mg^2+^ versus 10 µM+Mg^2+^)). (D) Representative current traces recorded at a membrane potential of -60 mV showing inhibitory kinetics of 10 μM K2060 at the GluN1/GluN2A receptor. The K2060 was applied for different periods, as indicated. (E) The average τ_off_ values are shown after applying 10 μM K2060 for the indicated period as shown in D (F_(3,22)_ = 0.574, p = 0.638, one-way ANOVA). (F) Representative current traces recorded at a membrane potential of -60 mV showing an inhibitory effect induced by 10 μM K2060 at GluN1/GluN2A receptor (as indicated) after its washout by ECS for 10 s or 100 s before eliciting the current responses by L-glutamate. (G) The current traces show the time course of the responses of the GluN1/GluN2A receptor after 10 s (red) and 100 s (blue) application of ECS, as shown in panel F. The traces were scaled and aligned as indicated. The dotted line represents 200 ms after the re-application of L-glutamate. (H) The bar graph shows the relative mean responses measured 200 ms after L-glutamate re-application as indicated in G. (t_(10)_ = -7.488, ***p = <0.001, Student’s t-test). (I) Representative current traces recorded at -60 mV showing inhibitory kinetics of 10 μM K2060 at the GluN1/GluN2A receptor. The current response of the GluN1/GluN2A receptor was recorded for 100 s after the termination of the application of 10 μM K2060 as indicated. (J) Representative current traces showing no “tail currents” after co-applying 1 mM L-glutamate and 10 or 30 μM K2060 at the GluN1/GluN2A receptor.

Next, we examined whether K2060 exhibits MCI. First, we used a classical MCI protocol on HEK293 cells expressing the GluN1/GluN2A receptor involving 30 s K2060 treatment without L-glutamate. ECS was applied for 1 s, and GluN1/GluN2A receptors were activated by 1 mM L-glutamate. The resulting current response elicited by L-glutamate was then compared to control current responses elicited after the previous absence of K2060 (Fig. 4A). We found that the minimum I_MCI_/I_control_ values were ∼0.8 for 1 µM K2060, ∼0.5 for 10 µM K2060 and ∼0.3 for 100 µM K2060 (Fig. 4B), whereas we previously obtained ∼0.5 for 100 µM memantine (Kolcheva et al., 2021; Kolcheva et al., 2023). We next asked whether the duration of application of K2060 alters MCI at GluN1/GluN2A receptors. We found that application of 10 µM K2060 for 1 s resulted in a minimum I_MC_I/I_contro_l value of ∼0.7, but extending the application time of K2060 to 3, 10, 30, or 100 s resulted in similar minimum I_MCI_/I_control_ values of ∼0.5 (Fig. 4C-D). Thus, applying K2060 lasting >3 s was sufficient to induce stable MCI at the GluN1/GluN2A receptor; to compare our data with previous MCI measurements with memantine, we continued to use 30 s of application of K2060. In the next phase, we evaluated how long it takes to eliminate MCI, i.e., how quickly K2060 is washed off the membrane. Therefore, we used the MCI protocol, but after co-applicating L-glutamate and K2060, we applied ECS for 1, 3, and 10 s (Fig. 4E). This experiment showed that the minimum I_MCI_/I_control_ value increased to ∼0.6 after 3 s of application of ECS compared to 1 s of application of ECS and that extending the application time of ECS to 10 s increased the minimum I_MCI_/I_control_ value to ∼0.9, indicating that K2060 is largely removed from the membrane after 10 s (Fig. 4F). Interestingly, the time constant for recovery from MCI (τ_recovery_; see Methods) ranged from ∼6-7 s for 100 µM K2060 (Fig. 4G,H) and 100 µM memantine (Glasgow et al., 2018; Kolcheva et al., 2021). Recent data have shown that the GluN2A-M630 amino acid residue lines the fenestration of memantine via MCI (Wilcox et al., 2022). Therefore, we next assessed whether GluN1/GluN2A receptors with the mutated GluN2A-M630 residue exhibit altered MCI induced by K2060 (Fig. 4I). Our data revealed no significant differences in the minimum I_MCI_/I_control_ ratio for 10 μM K2060 at GluN1/GluN2A-M630A and GluN1/GluN2A-M630W receptors. The measured IC_50_ values for mutant GluN1/GluN2A receptors at membrane voltage -60 mV were only slightly higher compared with wild-type GluN1/GluN2A receptor (GluN1/GluN2A-M630W: 2.70 ± 0.34 (n=12); GluN1/GluN2A-M630A: 1.25 ± 0.15 (n=7)). Thus, our experiments suggest that K2060 uses a different mechanism of fenestration during MCI compared with memantine.

**Fig. 4.**
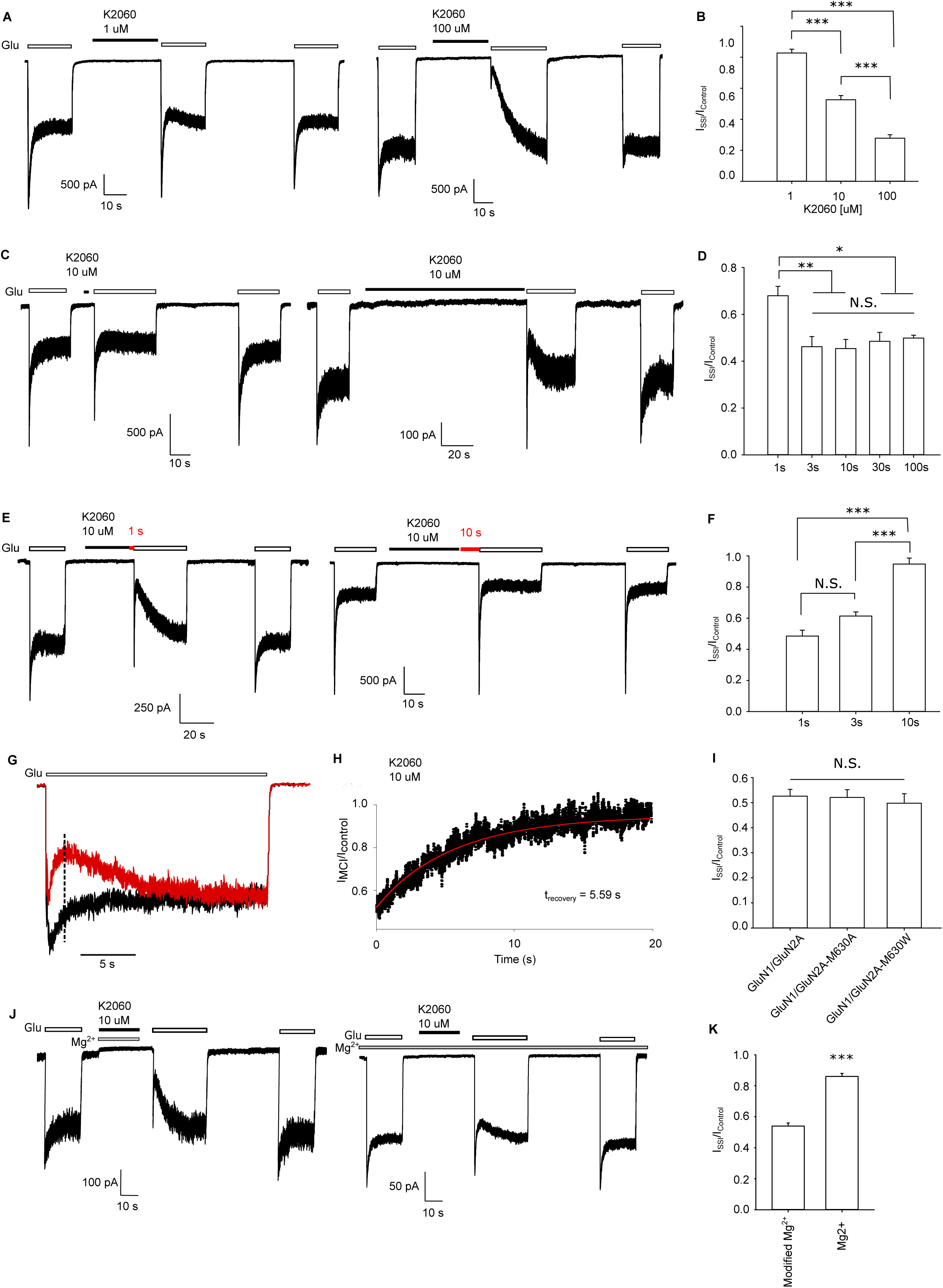
Membrane-to-channel inhibition induced by K2060. (A, C, E, J) Representative current traces from HEK293 cells expressing GluN1/GluN2A receptor showing different MCI protocols. (A) MCI protocol was induced by 1 µM or 100 µM K2060. (B) Summary of minimum I_MCI_/I_control_ values measured with the indicated concentrations of K2060 (1, 10, 100 μM) at GluN1/GluN2A receptors (F_(2, 17)_ = 70.433, ***p = <0.001, one-way ANOVA, Tukey’s post hoc test). (C) Adjusted MCI protocol was induced by different times of applying 10 µM K2060 (1 and 100 s). (D) Summary of minimum I_MCI_/I_control_ values measured after the different times of applying 10 µM K2060 (1, 3, 10, 30, and 100 s) (F_(4, 23)_ = 5.509, *p = <0.05,**p = <0.01 one-way ANOVA, Tukey’s post hoc test (versus 1 s of 10 µM K2060 application)). (E) Adjusted MCI protocol was induced by 10 µM K2060 with ECS washout times of 1 s or 10 s after the termination of the application of K2060; the ECS washout is marked in red. (F) Summary of minimum I_MCI_/I_control_ values measured with the indicated washout times (1, 3, and 10 s) described in C (F_(2, 16)_ = 35.426, ***p = <0.001 one-way ANOVA, Tukey’s post hoc test). (G) The representative control current trace induced by 1 mM L-glutamate (black) was aligned to the current trace obtained after applying 10 µM K2060 for 30 s (red; the standard MCI protocol). (H) I_MCI_/I_control_ was plotted from the example trace in panel C; the corresponding τ_recovery_ value is shown. (I) Summary of minimum I_MCI_/I_control_ values measured from HEK293 cells expressing GluN1/GluN2A, GluN1/GluN2A-M630A and GluN1/GluN2A-M630W receptors (F_(2, 19)_ = 0.215, p = 0.808, one-way ANOVA). (J) Adjusted MCI protocol was induced by 10 µM K2060 in the presence of 1 mM Mg^2+^ (”Modified Mg^2+^”), or 1 mM Mg^2+^ was present during the entire MCI protocol (”Mg^2+^”) as indicated. (K) Summary of minimum I_MCI_/I_control_ values measured in panel J. (t_(10)_ = -10.955, ***p = <0.001, Student’s t-test).

We next asked whether extracellular Mg^2+^ regulates K2060-mediated MCI. Therefore, we used the MCI protocol, i.e., co-applied 10 µM K2060 with 1 mM Mg^2+^ but employed the rest of the MCI protocol without Mg^2+^ (”modified Mg^2+^”; Fig. 4J). This experiment revealed a minimum I_MCI_/I_control_ value of ∼0.5, corresponding to the value measured for 10 µM K2060 in the absence of Mg^2+^ (Fig. 4K). Furthermore, we measured the entire MCI protocol in the presence of 1 mM Mg^2+^, and in this case, the minimum I_MCI_/I_control_ value increased to ∼0.9 (Fig. 4J,K). Our data showed a strong involvement of MCI in the mechanism of action of K2060 at the recombinant GluN1/GluN2A receptors.

To determine whether K2060 is a more potent inhibitor of native NMDARs than memantine, we measured evoked excitatory postsynaptic currents (eEPSCs) from CA1 neurons in acute slices from P30-35 aged mice (Fig. 5A-B). To simulate conditions from measurements of HEK293 cells and to mitigate the effects of intracellular Ca^2+^, we used 1 mM Ca^2+^ in the recording aCSF and 10 mM BAPTA in the ICS. We measured eEPSCs by stimulating Schaffer collateral connections from the CA3 region in the stratum radiatum with a concentric bipolar tungsten electrode. Stimulation consisted of 5 trains of extracellular stimuli at a frequency of 50 Hz under conditions of constant pharmacological inhibition of AMPARs and GABARs; then, we compared the amplitudes of the fifth response after 10 min of application of vehicle (DMSO) or memantine and K2060 (Fig. 5C). In the control experiments, we observed an average increase of ∼3%; for memantine, we observed a lower inhibition (∼21%) than K2060 (∼51%). The administration of memantine or K2060 showed no significant alteration in the decay time (tau of a single-exponential fit) of NMDAR eEPSCs. Therefore, these results showed that K2060 inhibited both recombinant and endogenous NMDARs more potently than memantine.

**Fig. 5.**
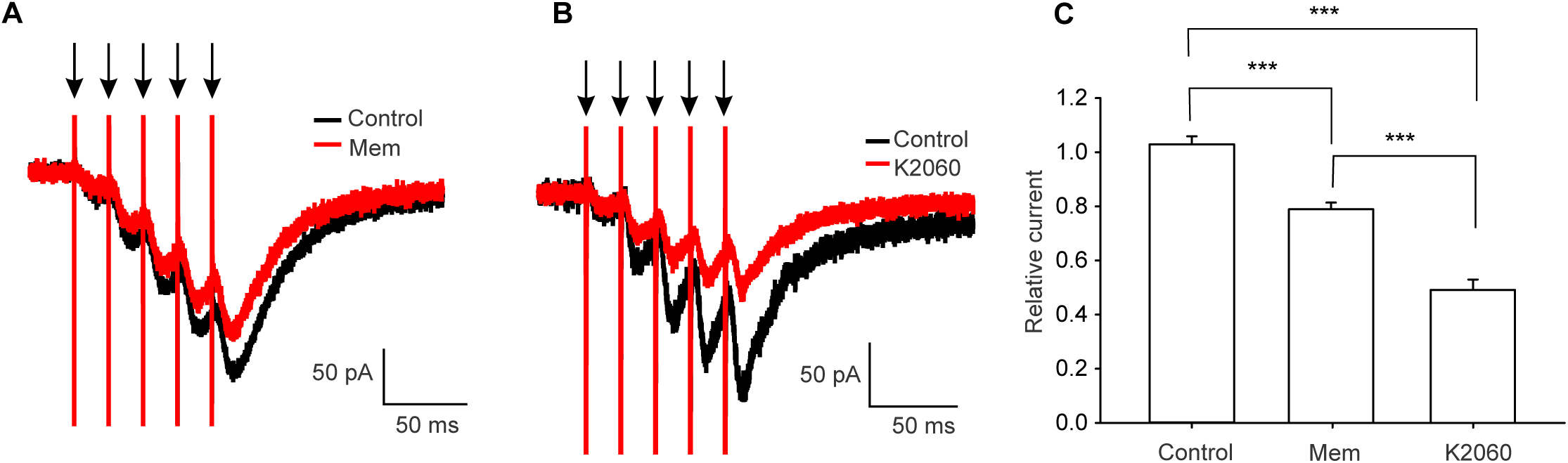
K2060 is a more potent inhibitor of endogenous synaptic NMDARs than memantine. (A, B) Representative averaged current traces showing NMDAR-EPSCs recorded from pyramidal CA1 neurons in control conditions (black traces) and after 10 min application of 30 µM memantine (A) or 30 µM K2060 (B) (red traces). NMDAR-EPSCs were evoked by trains of 5 extracellular stimuli (arrowheads) at 50 Hz. (C) Summary of relative current responses measured at the fifth response after 10 min treatment with DMSO (control; n = 5 cells from 3 mice), 30 µM memantine (Mem; n = 5 cells from 4 mice), and 30 µM K2060 (n = 5 cells from 3 mice) as indicated in panels A and B (F_(2,12)_ = 75.174, ***p = <0.001, one-way ANOVA, Tukey’s post hoc test). The administration of memantine or K2060 yielded no significant alterations in the decay time (tau of a single-exponential fit, p = 0.766, one-way ANOVA) of NMDAR eEPSCs.

### 3.3. In vivo effects of K2060

In assessing the potential of K2060 as a therapeutic compound targeting NMDARs, we evaluated its pharmacological attributes, including toxicity, pharmacokinetics, distribution, and neuroprotective capabilities in animal models. The initial step involved examining K2060’s acute toxicity in male and female mice by administering i.p. doses of 10, 15, 20, and 25 mg/kg following established protocols. Observations included sporadic increases in activity at doses of 10 mg/kg and 15 mg/kg, while doses of 20 mg/kg and 25 mg/kg induced pronounced symptoms such as whole-body tremors, impaired coordination, and disturbances in balance on one side, manifesting within 3-4 minutes post-administration. Despite the resolution of symptoms within two hours and the absence of adverse histopathological outcomes in the liver and kidneys, the No Observed Adverse Effect Level (NOAEL) was determined to be 15 mg/kg, with 20 mg/kg and 25 mg/kg as the maximum tolerated dose (MTD) in males and females, respectively. A more conservative dose of 10 mg/kg was selected for subsequent studies to ensure safety.

To determine the potential of K2060 as a drug for targeting the CNS, we examined its pharmacokinetic profile and brain distribution using a mouse model, with a 10 mg/kg dose administered intraperitoneally over 6 hours (Fig. 6). This specific dose achieves an adequate concentration (approximately 2 µM) within the brain, aligning with the measured inhibitory potency for the NMDARs under study. The K2060 was absorbed quickly, with peak concentrations (T_max_) reached at 5 minutes in plasma and 15 minutes in the brain, followed by prompt clearance from both compartments. The parallel peak concentrations (C_max_) observed in plasma and brain tissue suggest that the K2060’s entry into the brain occurs through passive diffusion.

**Figure 6.**
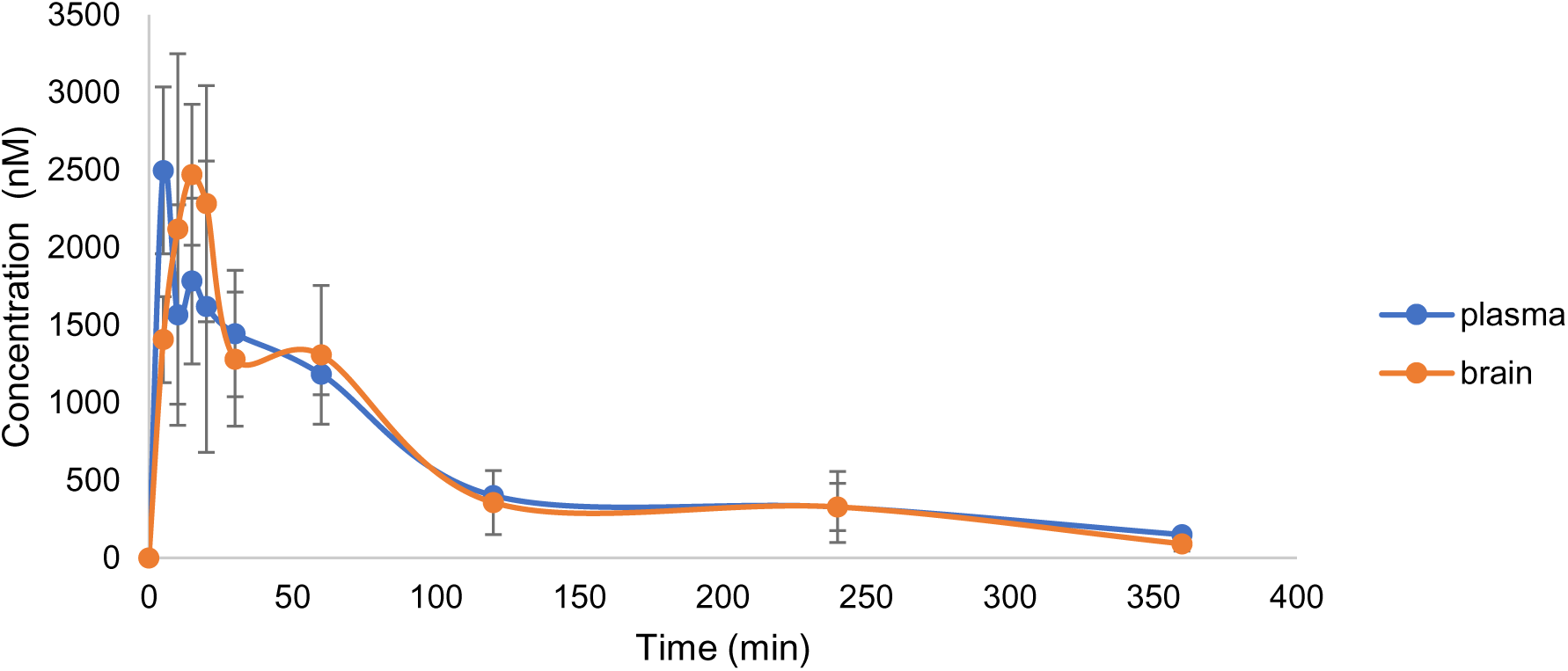
Pharmacokinetics of K2060 and its distribution to the brain. K2060 (10 mg/kg) was administered intraperitoneally to the male mice. Data are visualized as the mean ± S.E.M. (n = 4). Note that Tmax (plasma) = 5 minutes and Tmax (brain)=15 minutes in the brain, followed by prompt clearance from both compartments. Cmax (plasma)=2947 ± 537 nM; Cmax (brain)=2469 ±454 nM.

As the potential antidotal properties of NMDAR inhibitors are at the forefront of experimental treatments for organophosphorus poisoning (Gorecki et al., 2022), we further focused on a mouse model poisoned with tabun, a nerve agent that irreversibly blocks acetylcholinesterase (AChE). This blockage is notably resistant to reversal by oxime reactivators, making survival heavily reliant on the symptomatic relief provided by atropine and benzodiazepines. We explored the survival impact on mice using the K2060 as a standalone treatment and in combination with the established antidote regimen. The results revealed that K2060 alone did not offer any protective benefits. However, combined with the conventional antidote approach, specifically trimedoxime and atropine, significantly improved therapeutic outcomes were achieved (refer to Table 3).

## 4. Discussion

Open-channel blockers of NMDARs, such as memantine and ketamine, have been explored as potential therapeutics for various neurological and neuropsychiatric disorders. Memantine is approved for therapeutic use in AD (Robinson and Keating, 2006), and ketamine is approved for depression treatment (Kim et al., 2019; Mahase, 2019). MK-801 is another potent open-channel blocker of the NMDAR with high affinity and almost irreversible off-rate (Huettner and Bean, 1988). Initially explored for its anticonvulsant and anxiolytic effects (Sircar et al., 1987), its substantial side effects limit its clinical application (Janus et al., 2023). The contrast with well-tolerated memantine, which binds weakly, suggests the importance of kinetics and binding mechanisms for the clinical efficacy of the open-channel blockers of the NMDARs (Song et al., 2018). Memantine and ketamine, with their distinct electron distribution and structural features, highlight the complexity of NMDAR interaction. Specifically, K2060, unlike ketamine or memantine, is endowed with two aromatic rings that surround a central cycloheptenyl moiety, indicating a unique electron distribution that may influence its interaction with NMDARs. This structural aspect suggests the importance of hydrophobic interactions facilitated by aromatic rings, as these interactions are crucial for affinity towards NMDAR. The “irreversible” blockade of MK-801 is thought to be due to a robust hydrophobic interaction between its benzene rings and aromatic amino acid residues in the receptor pore (Lu et al., 2017; Song et al., 2018). Therefore, to avoid irreversible binding and the associated psychotomimetic side effects, most MK-801 analogs are synthesized as open or fully open-chain compounds with one or two cleaved benzene rings, respectively (Bachurin et al., 2001; Berger et al., 2009). Although K2060 has preserved benzene rings, it can still presumably leave the ion channel pore region, albeit with slower kinetics than memantine and ketamine (Kolcheva et al., 2023). This effect may be attributed to the cyclic alkyl moiety as well as exocyclic secondary amine moiety in the structure of K2060, which effectively constrains the molecule’s conformation without altering the chemical and physical properties of the original compound (Ono et al., 2002).

Our measurements without extracellular Mg^2+^ revealed that K2060 exhibited IC_50_ values between 0.6-0.7 µM at GluN1/GluN2A and GluN1/GluN2B receptors (Table 2). These values were more aligned with IC_50_ values for memantine and ketamine (1.8 µM and 0.9 µM for GluN1/GluN2A, 0.7 µM and 0.4 µM for GluN1/GluN2B, respectively) (Glasgow et al., 2017) than for (+)MK-801 IC_50_ values (0.013-0.3 µM and 0.009-0.07 µM for GluN1/GluN2A and GluN1/GluN2B, respectively) (Monaghan and Larsen, 1997; Dravid et al., 2007; Bettini et al., 2022). It has been shown that several open-channel blockers, including memantine and ketamine, compete for their binding site in the ion channel with Mg^2+^ at physiological levels (Kotermanski and Johnson, 2009; Glasgow et al., 2018; Kolcheva et al., 2023). In the presence of extracellular Mg^2+^, IC_50_ values for memantine increased ∼16-fold for GluN1/GluN2A receptors and ∼20-fold for GluN1/GluN2B receptors. Our measurements showed that the IC_50_ value for K2060 increased only ∼8-fold for GluN1/GluN2A receptors and ∼9-fold for GluN1/GluN2B receptors suggesting similar competing for Mg^2+^ as for memantine and ketamine. Furthermore, our experiments with K2060 in the presence of extracellular Mg^2+^ demonstrated analogous inhibition characteristics (τ_off_) and a reduction in MCI as observed in prior studies from memantine-induced inhibition in GluN1/GluN2A and GluN1/GluN2B receptors (Glasgow et al., 2018). In addition, Glasgow *et al*. showed that the time constant of MCI recovery (τ_recovery_) from memantine inhibition is consistent with the slow component of the recovery curve (τ_slow_) observed in the case of conventional memantine inhibition. Similarly, we observed a close overlap between τ_recovery_ and τ_slow_ in the case of K2060. In addition, we observed a decrease in τ_slow_ and A_slow_ in the presence of Mg^2+^, which may also be attributable to the reduction of MCI induced by K2060 in the presence of Mg^2+^. In the case of memantine, MCI was explained by the presence of a fraction of the uncharged form that interacts with the membrane and then fenestrates into the ion channel of the GluN1/GluN2 receptor (Wilcox et al., 2022). The stronger MCI of K2060 may be explained by differential lipophilicity (memantine clogP = 2.07; MK-801 clogP = 3.31; K2060 clogP = 4.59; MarvinSketch software v. 23.8; estimated values), although both memantine and K2060 are likely to wash off the membrane in the order of seconds (Kotermanski and Johnson, 2009). As MK-801 and K2060 are secondary amines, their basicity (MarvinSketch software v. 23.8 predicted values) also differs. MK-801 has a pKa value of 7.9, and the pKa for K2060 is predicted to be 8.8, meaning that under physiological conditions (pH = 7.3), only 34% of MK-801 but 97% of K2060 is protonated. Note that the pKa of K2060 is still within drug-like limits for centrally available compounds (Gupta et al., 2019). Interestingly, our measurements with NMDARs carrying mutations of the GluN2A-L630 residue showed that K2060 likely uses a different fenestration pathway than memantine (Wilcox et al., 2022), thus suggesting a different MCI mechanism than memantine.

**Table 2.** The inhibitory kinetics of K2060 at the GluN1/GluN2A receptor.

| Receptor | Mg<sup>2+</sup><br>(mM) | K2060<br>(uM) | $\tau_{on}$ (ms) | $\tau_{off}$ (ms) | $\tau_{off-fast}$ (ms) | A<sub>fast</sub> | $\tau_{off-slow}$ (ms) | A<sub>slow</sub> |
| --- | --- | --- | --- | --- | --- | --- | --- | --- |
| GluN1/<br>GluN2A | 0 | 3 | 1246.17 ±<br>175.78<br>(n=8) | 8629.05 ±<br>1361.69<br>(n=7) | 767.49 ±<br>361.73<br>(n=7) | 0.11 ±<br>0.04 (n=7) | 9571.44 ±<br>1550.35<br>(n=7) | 0.87 ±<br>0.04 (n=7) |
|  | 1 | 3 | 2727.82 ±<br>212.27<br>(n=8) | 4138.81 ±<br>335.01 (n=8) | 740.97 ±<br>104.35<br>(n=7) | 0.27 ±<br>0.05 (n=7) | 5416.06 ± 508.40<br>(n=7) | 0.73 ±<br>0.05 (n=7) |
|  | 0 | 10 | 384.92 ±<br>95.21 (n=9) | 8943.33 ±<br>1043.33<br>(n=9) | 829.47 ±<br>259.45<br>(n=9) | 0.10 ±<br>0.03 (n=9) | 9712.87 ±<br>1062.08 (n=9) | 0.90 ±<br>0.03 (n=9) |
|  | 1 | 10 | 1379.12 ±<br>110.18<br>(n=5) | 4494.98 ±<br>372.5 (n=5) | 806.33 ±<br>256.52<br>(n=5) | 0.18 ±<br>0.06 (n=5) | 5429.33 ± 777.85<br>(n=5) | 0.82 ±<br>0.06 (n=5) |

**Table 3.**
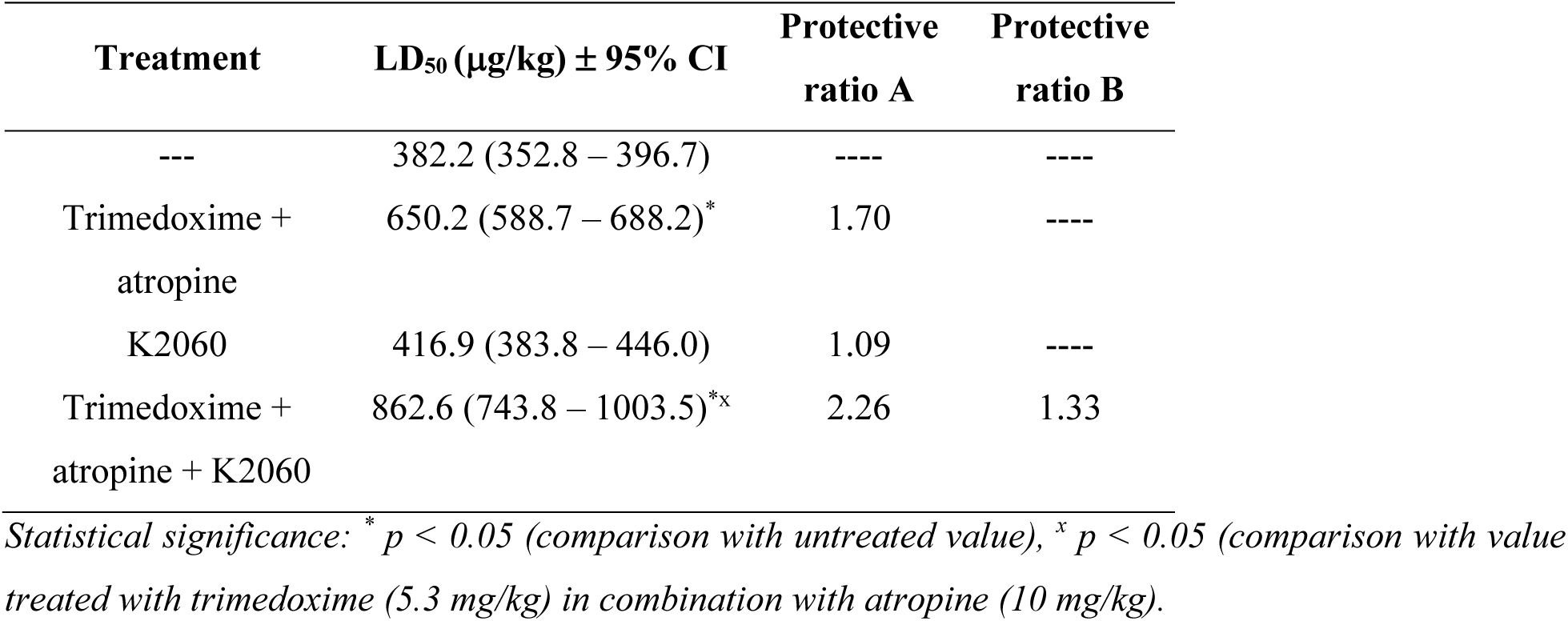
Therapeutic efficacy of K2060 (10mg/kg) in tabun-poisoned mice.

| Treatment | LD<sub>50</sub> (µg/kg) ± 95% CI | Protective ratio A | Protective ratio B |
| --- | --- | --- | --- |
| --- | 382.2 (352.8 – 396.7) | ---- | ---- |
| Trimedoxime +<br>atropine | 650.2 (588.7 – 688.2)* | 1.70 | ---- |
| K2060 | 416.9 (383.8 – 446.0) | 1.09 | ---- |
| Trimedoxime +<br>atropine + K2060 | 862.6 (743.8 – 1003.5)*<sup>x</sup> | 2.26 | 1.33 |

Statistical significance: \* $p < 0.05$ (comparison with untreated value), <sup>x</sup> $p < 0.05$ (comparison with value treated with trimedoxime (5.3 mg/kg) in combination with atropine (10 mg/kg)).

Previous studies using electrophysiological measurements in hippocampal slices have shown that memantine exhibits inhibition of synaptic NMDARs in CA1 pyramidal neurons only at >10 µM concentrations (Riebe et al., 2016). Our measurements on CA1 pyramidal neurons from hippocampal slices revealed a more pronounced inhibitory effect of 30 µM K2060 on NMDAR eEPSCs than 30 µM memantine. However, the extent of memantine-induced inhibition on pyramidal CA1 neurons at the same concentration was lower than previously reported using field excitatory postsynaptic potentials (fEPSP) (Riebe et al., 2016). The observed difference can be partially attributed to the technique and variations in the composition of recording solutions. This conclusion is consistent with a previous study that showed that inhibition of synaptic NMDARs by memantine in cortical pyramidal neurons is enhanced under conditions that promote intracellular Ca^2+^ accumulation (Glasgow et al., 2017). Our measurements in hippocampal slices were intentionally designed to mimic our recordings in HEK293 cells, thereby averting intracellular Ca^2+^ accumulation—a distinction from the fEPSP study in CA1, where such prevention was not implemented (Riebe et al., 2016). Moreover, our findings from hippocampal slices are consistent with our results from HEK293 cells, where we found more potent inhibition induced by K2060 compared to memantine at both GluN1/GluN2A and GluN1/GluN2B receptors. Consistent with previous publications on the inhibitory effect of memantine, we did not observe any alterations in the washout time constants of memantine or K2060 on NMDAR eEPSCs (Wild et al., 2013; Povysheva and Johnson, 2016).

In our *in vivo* studies, we sought to assess the drug-like properties of K2060 and the practical application of our *in vitro* findings on K2060. When given intraperitoneally, we determined that K2060 is relatively non-toxic, with a safe dosage threshold of up to 15 mg/kg after i.p. administration. This starkly contrasts the structurally related MK-801, which typically requires a dose of about 0.1 mg/kg to elicit effects on behavior, cognition, or social interaction (Nordquist et al., 2008; Vojtechova et al., 2023). Conversely, memantine is deemed safe at doses up to 20 mg/kg intraperitoneally, although doses between 5-10 mg/kg may induce some memory impairments in rats (Parsons et al., 1999). Regarding the distribution, K2060 was found to cross the blood-brain barrier effectively, with plasma and brain concentrations aligning closely, albeit with a 10-minute delay in peak concentration times in the brain. Despite the rapid clearance of the compound from both plasma and brain, concentrations of approximately 300 nM (about 13% of the peak concentration) were still detectable 4 hours post-administration in both compartments. Given the IC_50_ values for the GluN1/GluN2A and GluN1/GluN2B receptors (approximately 0.6-0.7 µM), a 10 mg/kg dose proved sufficient concentrations above 1 µM even an hour after dosing. Finally, as the NMDAR antagonists are recognized for their positive effects in protecting against the severe seizures triggered by organophosphates (Figueiredo et al., 2023; Pulkrabkova et al., 2023), we explored K2060’s impact on survival rates in mice subjected to tabun exposure. We found that including K2060 in conventional antidote treatments (atropine+trimedoxime) significantly enhanced the LD_50_ value and overall survival rates. In comparison, pretreatment with memantine alone or even supplemented by HI-6 and atropine after exposure to soman did not cause significantly better survival (Kassa and Zdarova Karasova, 2023). Other studies have shown some beneficial effects with memantine pretreatment (Stojiljkovic et al., 2019), indicating that certain doses of memantine might have a partial effect (Dai et al., 2004). Thus, K2060 showed promising results against severe organophosphate-induced seizures and could represent a new class of NMDAR inhibitors with unique inhibitory properties that could find application in other pathologies associated with the dysregulation of NMDARs.

## CRediT author statement

**Anna Misiachna**: Conceptualization; Data curation; Formal analysis; Investigation; Visualization; Roles/Writing - original draft; **Jan Konecny**: Investigation; Methodology; Writing - original draft; **Marharyta Kolcheva**: Conceptualization; Data curation; Formal analysis; Investigation; Methodology; Roles/Writing - original draft; **Marek Ladislav**: Conceptualization; Data curation; Formal analysis; Investigation; Methodology; Roles/Writing - original draft; **Lukas Prchal**: Investigation; Methodology; Data curation; Writing - original draft; **Jakub Netolicky**: Data curation; Formal analysis; **Stepan Kortus**: Formal analysis; **Petra Barackova**: Investigation; Methodology; **Emily Langore**: Methodology; Resources; **Martin Novak**: Investigation; Methodology; **Katarina Hemelikova**: Investigation; Methodology; **Zuzana Hermanova**: Investigation; **Michaela Hrochova:** Investigation; **Anezka Pelikanova:** Investigation; **Jitka Odvarkova**: Investigation; Methodology; **Jaroslav Pejchal**: Investigation; Methodology; **Jiri Kassa**: Investigation; Methodology; Funding acquisition, Writing - original draft; **Jana Zdarova Karasova**: Investigation; Methodology; Funding acquisition, Writing - original draft;; **Jan Korabecny**: Investigation; Methodology; Funding acquisition, Writing - original draft;; Supervision; Validation; Roles/Writing - original draft; **Ondrej Soukup**: Conceptualization; Data curation; Funding acquisition; Project administration; Resources; Supervision; Roles/Writing - original draft; **Martin Horak**: Conceptualization; Data curation; Funding acquisition; Project administration; Resources; Supervision; Roles/Writing - original draft.

## Acknowledgments

This work was supported by the Czech Science Foundation (24-10026S), project registration number CZ.02.01.01/00/22_008/0004562 (ExRegMed, MEYS CR), and the Ministry of Defence of the Czech Republic “Long Term Organization Development Plan 1011”—Healthcare Challenges of WMD II of the Military Faculty of Medicine Hradec Kralove, University of Defence, Czech Republic (Project No: DZRO-FVZ22-ZHN II).

## Data availability

Data will be made available on request.

## Abbreviations

AChE, acetylcholinesterase; AD, Alzheimer’s disease; AMPAR, α-amino-3-hydroxy-5-methyl-4-isoxazole propionic acid receptor; aCSF, artificial cerebrospinal fluid; CNS, central nervous system; DMSO, dimethyl sulfoxide; eEPSCs, evoked excitatory postsynaptic currents; ECS, extracellular recording solution; FBS, fetal bovine serum; fEPSP, field excitatory postsynaptic potential; GABAR, gamma-aminobutyric acid receptor; HEK293, human embryonic kidney 293; IC_50_, half-maximal inhibitory concentration; ICS, intracellular recording solution; IS, internal standard; LD_50_, median lethal dose; 7-MEOTA, 7*-*methoxy derivative of tacrine; MCI, membrane-to-channel inhibition; NMDAR, *N*-methyl-D-aspartate receptors; NOAEL, No Observed Adverse Effect Level; PBS, phosphate-buffered saline; RT, room temperature; TMD, transmembrane domain.

